# EpiState-T for representing epigenetic states across time

**DOI:** 10.64898/2026.11.12.766055

**Authors:** Jade Sunghyeok Choi, Jung In Choi

## Abstract

Epigenetic information is central to cellular identity, aging and rejuvenation. Unlike the relatively stable genomic sequence shared by most somatic cells, epigenetic states vary across cells and time, making their dynamics difficult to reconstruct from sparse experimental measurements. We introduce EpiState-T, the Temporal Epigenetic State Tensor, to represent selected epigenetic states across intervals and distinguish an observed trajectory from the use of a retained record. In a ten-cell simulation, an explicit reader can act on each cell’s surviving baseline record. Across 500 paired simulations, mean final distance to the cell’s own baseline was 0.736 with reading disabled, 0.182 with correctly assigned records and 0.402 with reassigned records. These differences follow from the specified reader. We separately applied the observation and transition tensors to 859,151 CpGs measured in three donor-associated fibroblast cultures before, during and after transient reprogramming. Among sites ending within 0.05 methylation units of baseline, 9.75–15.72% had first departed in treated cultures, compared with 5.37–9.91% in matched mock cultures. The intermediate measurement therefore distinguishes paths that an endpoint comparison merges. These are paths of population measurements: the human data do not track individual cells or establish that cells read a historical record. EpiState-T provides an initial mathematical framework for investigating temporal epigenetic information while keeping state representation, observed change and proposed biological mechanisms distinct.

## 1 Introduction

Understanding cellular aging and rejuvenation requires investigating how biological information is maintained and modified over time. Although most somatic cells share a largely stable genomic sequence, their epigenetic states vary across cell types and individual cells and change in response to developmental processes, aging and environmental conditions. Variation across cells and over time makes a cell’s history difficult to reconstruct from a few measurements. For example, a culture’s average methylation at a site may end near its starting value whether it stayed near that value or moved away and returned. An intermediate measurement can reveal a departure, but a culture average cannot tell us whether the same cells returned to their earlier states. We developed EpiState-T to make explicit both the path the measurements reveal and the biological questions they leave open.

A molecular profile measured after an intervention can resemble an earlier profile for several reasons. Present regulation can converge on a familiar state, some features can persist throughout treatment, or features can depart and later return. A further hypothesis proposes that retained historical information directs the later response. Endpoint resemblance alone cannot distinguish these descriptions. Their separation requires explicit time indexing, a definition of retained content and a testable account of how that content affects subsequent transitions.

Transient reprogramming provides an accessible setting for this measurement problem. Gill and colleagues released molecular profiles collected before, during and after a defined intervention in human fibroblasts [1]. An age-associated predictor, exemplified by methylation clocks [2], addresses a different target from resemblance to a particular pretreatment sample. Neither target by itself identifies retrieval of the cell’s own earlier state. The present work concerns the representation and analysis needed to keep these questions distinct.

Tensors already organize multiway data [3]. Tensor Composition Analysis estimates cell-type-specific methylation signals from bulk profiles [4], EpiTensor integrates epigenomic measurements to infer chromatin interactions [5], and kernel tensor methods can analyze multiple molecular data types with shared experimental indices [6]. The contribution of EpiState-T is a specification for identifying the trajectory unit, observation times and measured features, then distinguishing these observations from candidate interactions and retained content. It does not require a new tensor decomposition.

Jung In Choi proposed temporal links and preservation of past epigenetic information by analogy to properties of distributed ledgers [7,8]. Here we formalize two questions motivated by that proposal: what can an observer reconstruct from samples collected at known times, and what changes if a cell can use a retained record? A link in the research record identifies prior observations. It does not imply that cells hash, authenticate or preserve earlier states unchanged.

Section 2 defines the representation and the changes that can be reconstructed from successive measurements. Section 3 implements a reader in virtual cells, where both the retained record and access to it are known. Section 4 constructs observation and transition tensors from human fibroblast measurements, works through one measured site, and applies the same return rule to matched mock cultures. In the human data neither a cell-to-cell graph nor a reader is observed.

## 2 The Temporal Epigenetic State Tensor

### 2.1 Units, intervals and measurements

Let u identify a trajectory unit, p a molecular feature and k an interval with boundary times *t*_0_ ,*…, t* _*K*_. Unit type is mandatory metadata. It can identify a tracked cell in a simulation or a donor-associated population in a bulk experiment; the latter must never be relabeled as repeated observations of the same cell. Within an implementation, identifiers, feature order and interval conventions remain explicit. We use *I* _*k*_ =[ *t*_*k* −1_ , *t*_*k*_) with the final endpoint retained as a separate boundary observation. In the jump identity below, boundary states are taken immediately before a boundary event. A jump at the left endpoint is therefore assigned to the interval that follows it. The simulation has no molecular jumps exactly at its boundaries.

Table 1 summarizes the specification. Unmeasured components remain explicitly unavailable. Constructing an observation array does not imply that every hypothesized mechanism has been measured.

**Table 1.** What is known in the simulation and what is measured in the fibroblast data.

| Object | Meaning | Synthetic example and public-data application |
|---|---|---|
| M and optional R | Modeled molecular and spatial states | Known simulator states; no inferred latent cellular states or positions in the public case |
| O and Q | Observations and availability | Complete synthetic observations; QC-passed bulk beta measurements in the public case |
| G | Directed sender-to-recipient influence weights | Assumed spatial coupling in simulation; unavailable in the public case |
| E and Δ | Complete events or observed increments | Complete simulator jumps; two observed phase differences in the public case |
| H and Read | Retained content and its functional use | Explicit local checkpoint and reader in simulation; unavailable in the public case |
| Metadata | IDs, units, times, provenance and versions | Declared in both examples; supports reproducibility without authenticating biology |

For modeled molecular states and available positions, define

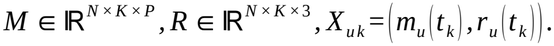

The baseline at *t*_0_ is stored separately. Molecular features and physical coordinates retain their own units; concatenation does not justify an unscaled distance mixing them. Physical x, y and z are feature coordinates, rather than the three modes of a cell-by-interval-by-feature tensor.

Observations are distinguished from modeled states:

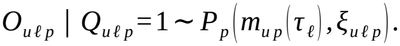

Here Q records assay availability, *τ* _*ℓ*_ is the observation time or declared protocol phase, and *ξ* collects measurement conditions. Missing observations are unavailable values, not methylation zeros. A methylation-array beta value is an observed intensity-derived fraction estimate; it is not an exact count of two DNA copies. The assay model and sampling unit must accompany the array.

### 2.2 Temporal increments and unresolved paths

For a completely observed piecewise-constant trajectory, signed molecular jumps satisfy

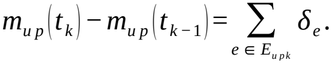

An empirical difference generally does not identify the underlying events. Between two endpoints a mark can remain constant or undergo a reversible excursion. Intermediate observations resolve some such ambiguity, but can still miss changes between samples. When the observations follow successive indexed phases, we store observed increments *Δ*_*uk p*_ = *O*_*uk p*_ *−O*_*u,k* −1, *p*_ without treating them as complete event logs. Unequal or incompletely specified time gaps prevent interpretation of these increments as rates unless actual durations are supplied.

The distinction is operational. An endpoint analysis asks where a trajectory ends. A path analysis additionally asks whether it changed direction, departed temporarily, or changed only in a later sampled interval. These comparisons can be computed from the observation array even when endogenous historical storage is unmeasured.

### 2.3 Selective retention and functional access

Let *H*_*u,k* −1_ denote previously active content, *Π*_*uk*_ a retention operator and *U*_*uk*_ newly encoded content:

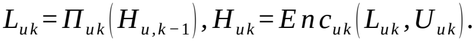

The operator may preserve, transform or lose information. Source time and present encoding time are distinct. Re-storage means forwarding selected prior content while adding new content; it does not mean replacing an old value by the latest measurement. A physical carrier need not occupy a separate compartment, and historical content could overlap current regulatory state. Here L denotes the portion carried forward.

Retention alone does not establish functional memory. A candidate transition law makes the additional reading hypothesis explicit:

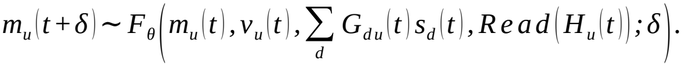

The input v denotes current external or experimental conditions, and s denotes current sender signals. The no-reading comparator disables Read while retaining the current state, measured inputs and appropriate model capacity. A flexible latent H can absorb unmeasured persistent variables; improved prediction alone would not identify a biological archive. The reader in Section 3 is a fully specified synthetic mechanism rather than an inferred molecular process.

## 3 Executable example with functional historical reading

### 3.1 Background dynamics and initial conditions

The example has ten fixed cell identifiers, three fictitious CpGs, two ideal binary copies per feature, and five intervals with boundaries 0 through 5 in arbitrary model time. Let j index the two copies. Define

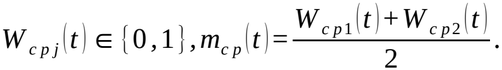

This idealization permits fractions 0, 0.5 and 1. It does not represent strand-resolved hemimethylation or the measurement model of the public array. The endpoint tensor has shape 10 *×* 5 *×* 6, comprising three fractions and three model coordinates; separate M and R arrays each have shape 10 *×* 5 *×* 3.

At each interval start, the symmetric union of three-nearest-neighbor relations defines the assumed graph. Pair weights are exp[ − (*d* / 1.4)^2^] with zero diagonal. Positions follow prescribed sinusoidal displacements; the graph weights are calculated at the start of each interval and held fixed until the next boundary. The synthetic display index and normalized sender input are

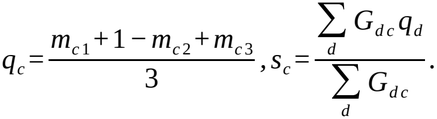

An isolated receiver has input zero. The index q is not a trained aging clock. Eligible background copy flips occur at rate

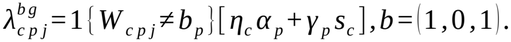

The vectors *α* = (0.120 , 0.100 , 0.110) and *γ* = (0.42 , 0.36 , 0.40) are illustrative. Cell factors *η*_*c*_ are evenly spaced from 0.85 to 1.15. Initial methylated-copy counts across the twenty copies are 2, 16 and 4 for the respective CpGs, assigned by the simulation seed. The background drive continues after reading is enabled.

### 3.2 Reading a surviving local record

At t=0 each cell encodes one self-contained baseline payload containing its three molecular fractions. Event records are appended at boundaries. Previously active records independently survive a boundary with probability ρ before new records are added. Retention draws are keyed by seed, boundary, cell and record identifier, keeping baseline-survival draws matched across arms despite different numbers of new events.

The declared query selects the baseline checkpoint at source time *t*^*⋆*^ = 0 from the cell’s currently active H. It returns a target 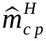 and availability indicator *χ*_*c p*_. If the checkpoint is lost, or access is blocked, *χ*_*c p*_ = 0. New event records cannot recreate a missing baseline. The transition routine cannot read the evaluator’s baseline array, an external registry or future records. Accurate encoding and source-time addressing are explicit engineering assumptions.

Let [ *x*]_+_ = *max* (*x* , 0) and define

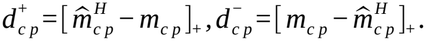

The added channel has rate

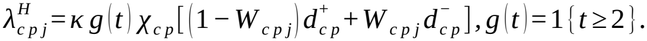

The total copy-flip rate is *λ*^*b g*^ + *λ*^*H*^. Reading permits upward flips when a fraction is below its retained target and downward flips when it is above. Its rate vanishes at the target, with unavailable content, with blocked access or when κ=0. The main active reading strength is κ=4. All rates are recomputed after each event and at interval boundaries. The first two intervals use background dynamics alone; reading begins after retention at the t=2 boundary. No state is externally reset to produce recovery. Figure 1 shows how the state tensor and baseline-distance trajectories change once reading starts.

**Figure 1.**
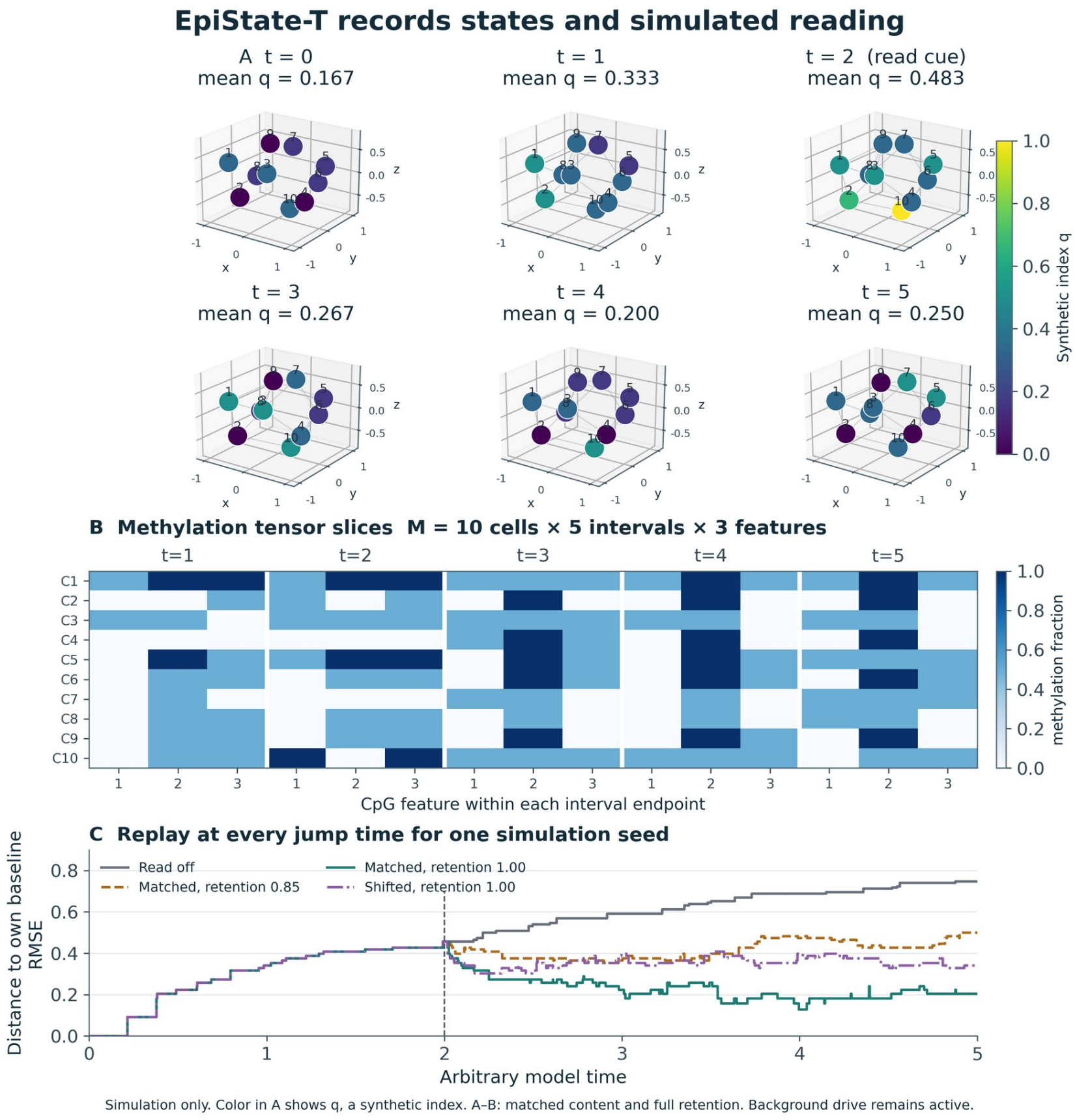
A simulated cell can use only a surviving record. (A) The same ten cells at baseline and at five interval endpoints; color represents the synthetic index q, not age. (B) Each slice shows three methylation features for all ten cells. A and B use matched records with full retention. (C) Exact event-time replay for four reading conditions with the same seed (20261106). Reading is enabled at t = 2; the background drive continues. Distance measures departure from each cell’s own baseline. No external reset produces the reversal.

### 3.3 Comparators and verification

The primary outcome measures distance to each cell’s own pre-change profile:

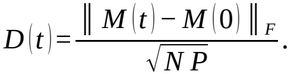

The Frobenius norm is taken over the cell-by-feature matrix. The evaluator uses the reference only for scoring. Six conditions compare reading off, matched reading at full or 0.85 retention, zero retention of prior records, access blockade with preserved records, and shuffled full-retention payloads. A fixed cyclic reassignment shuffles targets among cells while preserving present states and the population-level payload multiset. Content specificity is therefore compared between matched and shuffled arms at the same retention.

The primary experiment uses 500 independent seeds from 20261106 through 20261605, paired across conditions. Sensitivity calculations use the first 100 seeds for each combination of *ρ∈* {0 , 0.25 , 0.5 , 0.75 , 0.85 , 1} and *κ* ∈ {0 , 1 , 2 , 4 , 8}. Parameters were not optimized against a biological outcome. Intervals around means are 1.96 Monte Carlo standard errors across simulated populations, not uncertainty intervals for biological effects. The numerical grid is supplied in Supplementary Table S1.

Fourteen executable checks test numerical reproducibility, transition direction, replay of the event history, retention and loss of records, exclusion of missing or future content, the time at which reading starts and conditions under which reading should have no effect.

### 3.4 The effect of reading depends on the record

Mean final RMSE was 0.7355 with reading off, 0.1825 with matched full-retention reading and 0.4338 with retention 0.85 (Table 2; Figures 1 and 2). Zero retention and access blockade produced the same state paths as reading off in every replicate. Shuffled full-retention content gave RMSE 0.4025; the paired advantage of matched content was 0.2200, with a Monte Carlo interval of 0.2141–0.2258.

**Table 2.** Synthetic final baseline-distance RMSE over 500 paired replicates. Retention is per boundary. Intervals quantify Monte Carlo sampling error under the assumed mechanism.

| **Condition** | **ρ** | **Mean RMSE** | **95% Monte Carlo interval** |
|---|---|---|---|
| Reading off | 0.85 | 0.7355 | 0.7309–0.7401 |
| Matched local reading | 1.00 | 0.1825 | 0.1781–0.1868 |
| Matched local reading | 0.85 | 0.4338 | 0.4254–0.4421 |
| Zero retention of prior records | 0.00 | 0.7355 | 0.7309–0.7401 |
| Records retained but access blocked | 1.00 | 0.7355 | 0.7309–0.7401 |
| Shuffled local payloads | 1.00 | 0.4025 | 0.3982–0.4067 |

**Figure 2.**
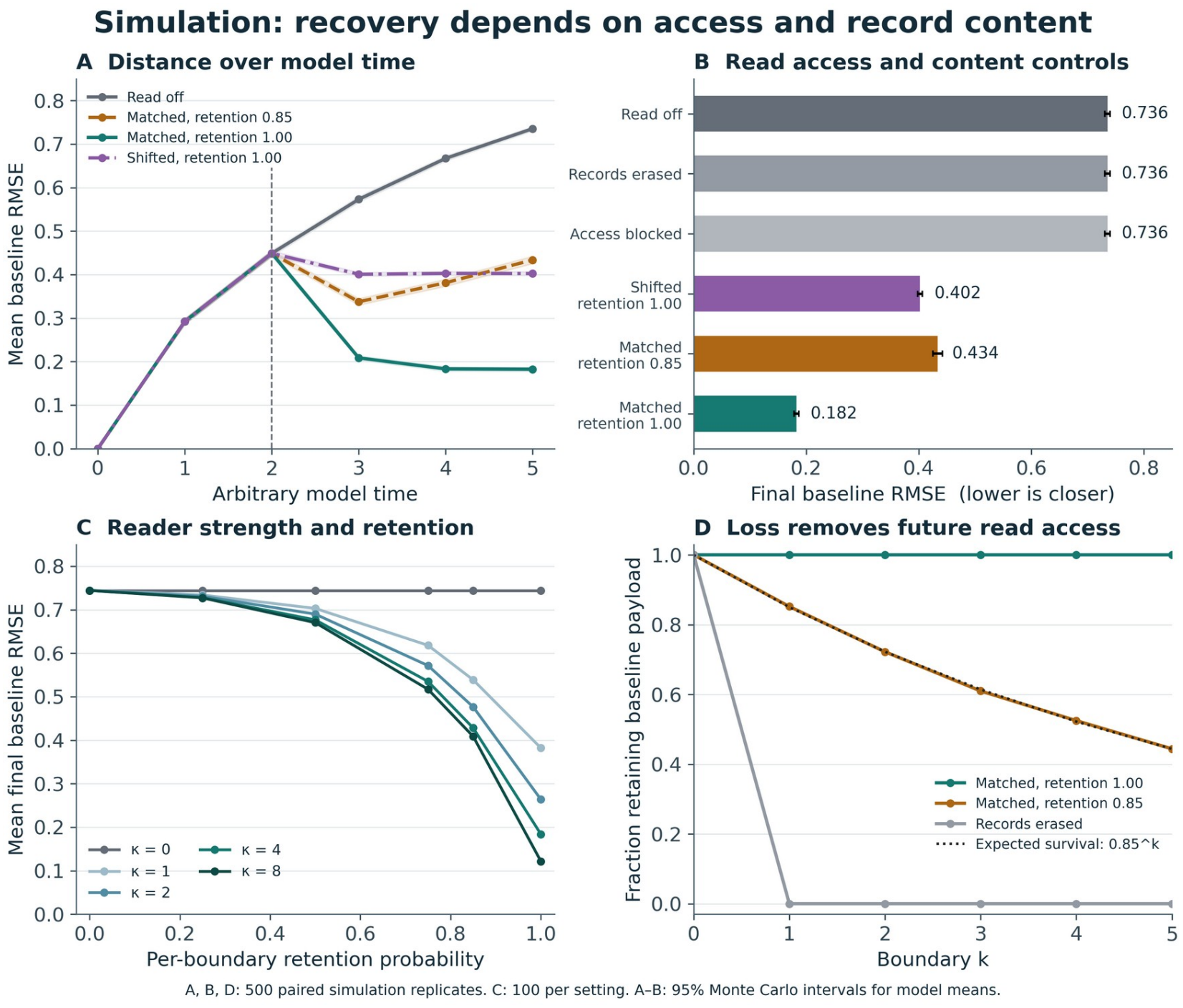
Correctly assigned records reduce final distance more than reassigned records in the simulator. (A) Mean distance to baseline over time. (B) Final distances, including erasure and blocked access. (C) Sensitivity to retention and reading strength (100 simulated populations per setting). (D) Survival of baseline records across intervals; the dotted line gives expected survival 0.85^*k*^ . A, B and D summarize 500 simulated populations. Uncertainty intervals describe simulation sampling error only.

Partial retention did not guarantee durable recovery. Mean RMSE fell from 0.4489 at t=2 to 0.3375 at t=3, then increased to 0.4338 at t=5 as records disappeared under continuing background drive. Shuffled content still improved over no reading because cells shared baseline features. Generic improvement therefore did not establish recipient-specific guidance.

The realization shown in Figure 1 uses full retention and seed 20261106 and contains 83 events, including 39 reading events, and 93 final records. The 0.85-retention realization has 72 events, 27 reading events and 63 records. These counts describe one realization. The intervention on the reader changes simulated trajectories; the direction of the response is specified by the model.

## 4 Observed methylation paths in human fibroblasts

### 4.1 Samples and preprocessing

We selected the methylation-array release GSE165179 from the Gill study within GSE165180 [1,9]. The ten-day exp2 branch supplies starting, intermediate and post-withdrawal profiles for donors O1, O2 and O3. The source experiment used adult dermal fibroblasts; the later phase followed several weeks without factor induction. These are destructively assayed donor-associated populations, rather than repeatedly measured individual cells. The baseline is an adult pretreatment culture and not a sample of that donor’s youthful state.

We used the archived, quality-filtered tensor from the processed-data analysis. Its preprocessing script and source-file SHA256 are retained in the accompanying package. The deposited matrix was reported as NOOB-normalized. The inherited filter retained cg probes with finite beta in [0,1] and finite detection P≤0.01 across the 23 profiles selected in that preprocessing: nine focal profiles, twelve control or failed-reprogramming profiles and two reference profiles. It retained 859,151 of 865,859 rows. This is the primary set. We separately assessed the effect of excluding flagged probes (§4.5); we did not reprocess raw IDAT files.

We recomputed the increment, path-class and backtracking results from the archived beta values. This calculation starts from the processed tensor, not from raw array files. To preserve the provenance of each measurement [10], the accompanying code records the source accession, sample mappings, input hashes and preprocessing steps.

### 4.2 Constructing the observation and transition arrays

For this dataset the observation array is

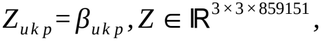

with modes donor-associated population, observed phase and CpG. The phases k=0,1,2 denote baseline B, intermediate I and after withdrawal A. They define two observed transition segments and are not five simulated intervals or equally spaced times. The QC-passed mask is complete for these retained features. Coordinates, intercellular edges, endogenous H and complete molecular events are unavailable and are not imputed. Sample linkage identifies donor and protocol condition only.

We apply EpiState-T by constructing the ordered transition tensor *Δ∈* ℝ^3*×*2*×*859151^ and maintaining feature IDs and source profiles:

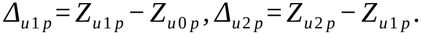

The closure *Δ*_*u*1 *p*_ + *Δ*_*u*2 *p*_ = *Z*_*u*2 *p*_ *−Z*_*u*0 *p*_ is checked for every entry. This step connects the representation to explicit path operators, rather than using the tensor only as a label for endpoint data. Closure is an arithmetic consistency check on indexing and subtraction; it supplies no independent evidence about the path between visits.

For example, at cg06036945 in donor O2 the treated culture measured 0.097, 0.312 and 0.122. The two entries in its transition tensor are approximately +0.216 and −0.190 beta units, calculated before rounding. Their sum, +0.026, records the small net change but hides the much larger excursion. The matched mock series was 0.097, 0.102 and 0.123: almost the same starting and final values, with no measured departure beyond 0.05. The probe passed both sensitivity filters. We chose this example from a fixed display panel by its treated trajectory before inspecting its mock values; it illustrates the calculation, not its prevalence (Supplementary Figure S1).

### 4.3 Measuring departures and returns

Write *a*_*u p*_ = *Δ*_*u*1 *p*_ and *r*_*u p*_ = *Z*_*u*2 *p*_ *−Z*_*u*0 *p*_. The sampled variation and backtracking are

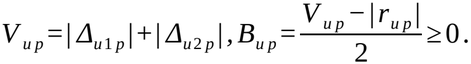

Backtracking equals *min* (| *Δ*_*u* 1 *p*_ | , | *Δ*_*u* 2 *p*_ |) when the increments oppose and is zero otherwise. It quantifies cancellation across the observed segments, not all unobserved changes between assays. Two profiles with the same baseline-to-final displacement can have different V and B.

Using an explicitly chosen tolerance ε=0.05 beta units, we classify each feature into six exhaustive categories. Near-baseline features satisfy | *a* | *≤ ε* and | *r* | *≤ ε*. Delayed departures satisfy | *a* | *≤ ε* but | *r* |> *ε*. Returns after departure satisfy | *a* |> *ε* and | *r* | *≤ ε*. Among features with both magnitudes above ε, same-sign a and r identify partial reversal when | *r* |<| *a* | and persistent or further departure otherwise; opposite signs identify a crossed baseline. These are descriptive observed-path labels, not molecular memory classes or statistical significance calls.

The tolerance was specified for this exploratory analysis, without calibration to assay error or a biological response. Sensitivity results use ε=0.02 and 0.10. Primary summaries use the full retained feature set; §4.5 reports the filtered sensitivity analysis. The heatmap in Figure 3 uses 120 randomly selected CpGs with seed 20261110, ordered by O1 baseline beta for display; no outcome-based feature selection enters the metrics. CpGs are not treated as independent donor replicates, and no feature-level significance test is used to support a population claim.

**Figure 3.**
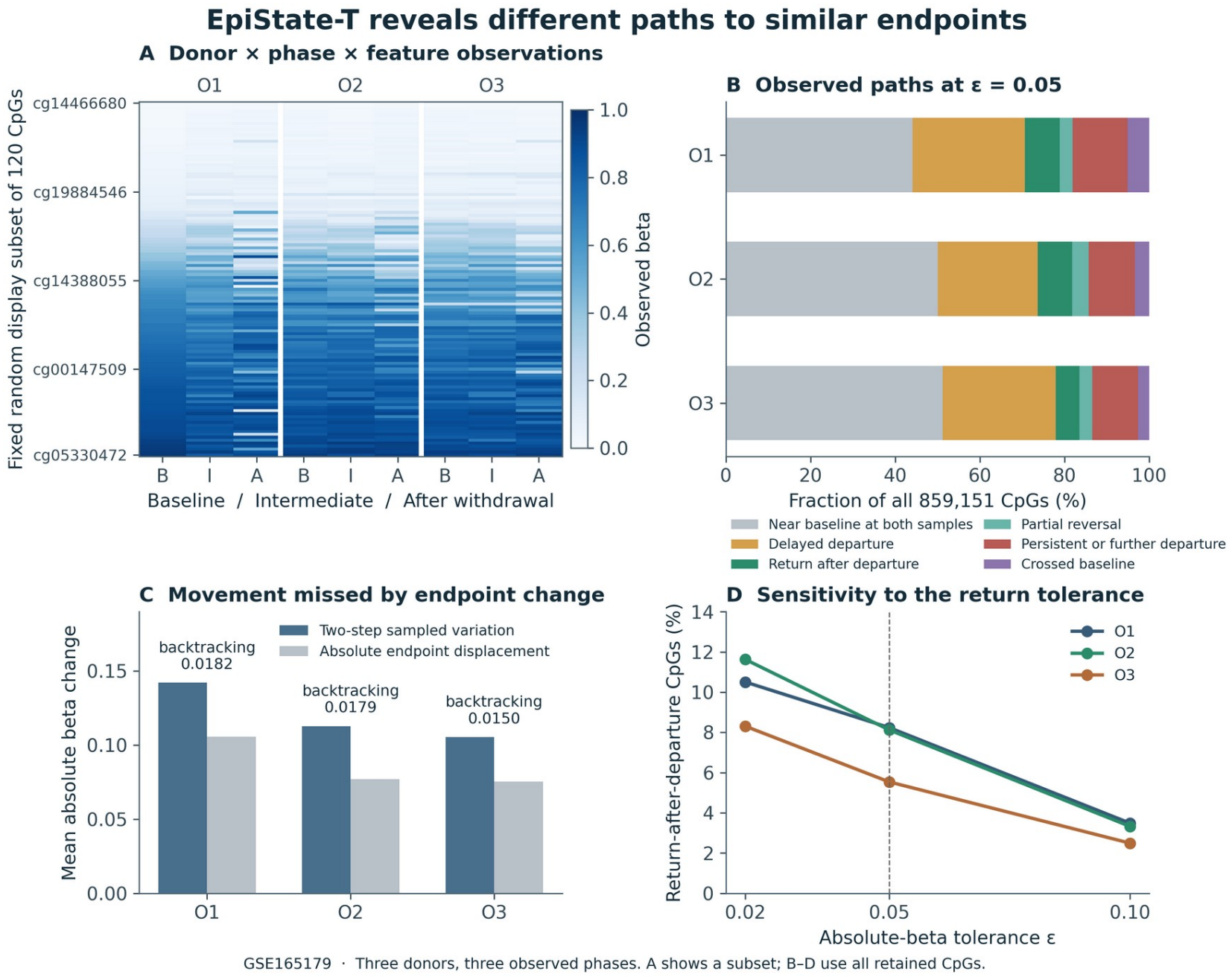
Similar endpoints conceal different sampled methylation paths. (A) A fixed display sample of 120 CpGs from the donor-by-phase array; all numerical summaries use 859,151 CpGs. (B) Six observed path classes at a tolerance of 0.05 beta units. (C) Sampled variation and absolute endpoint displacement; half their difference is the mean backtracking. (D) Fraction returning after departure at three tolerances. B, I and A denote baseline, intermediate and after withdrawal. Rows follow donor populations, not individual cells.

### 4.4 Sites with similar endpoints can follow different sampled paths

Whole-profile RMSE to baseline increased between I and A in all donors: 0.07972 to 0.16935 in O1, 0.07619 to 0.12473 in O2, and 0.07086 to 0.12025 in O3 (Table 3). Nonetheless, 70,733, 69,734 and 47,543 loci respectively met the observed return-after-departure definition: 8.23%, 8.12% and 5.53% of all retained features at ε=0.05.

**Table 3.**
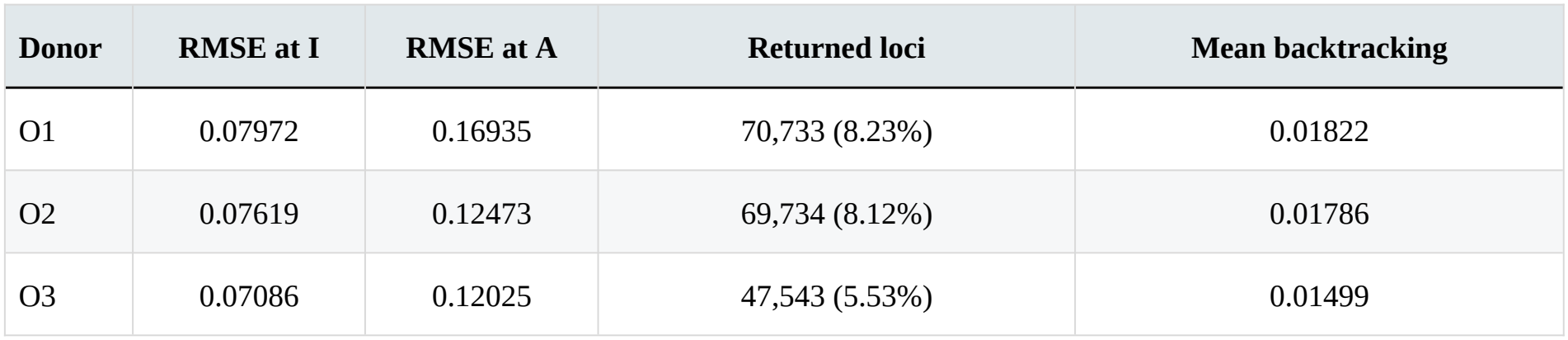
Changes in whole-profile distance and returns at individual CpGs by donor. Returns require an intermediate departure greater than 0.05 and a final deviation no greater than 0.05. Backtracking is the mean observed cancellation in beta units.

| Donor | RMSE at I | RMSE at A | Returned loci | Mean backtracking |
|---|---|---|---|---|
| O1 | 0.07972 | 0.16935 | 70,733 (8.23%) | 0.01822 |
| O2 | 0.07619 | 0.12473 | 69,734 (8.12%) | 0.01786 |
| O3 | 0.07086 | 0.12025 | 47,543 (5.53%) | 0.01499 |

Among features near baseline at the final phase, 15.72%, 13.95% and 9.75% had an observed intermediate departure. A baseline-to-final comparison would group these loci with those near baseline at both sampled phases. The intermediate measurement separates sites that otherwise have the same endpoint classification. This distinction does not require assuming that either class contains a biological archive.

Mean sampled variation exceeded absolute endpoint displacement in every donor. Mean backtracking ranged from 0.01499 to 0.01822 beta units. The global increment cosines were −0.0270, −0.1370 and −0.1209: some directional reversal can coexist with a larger final distance when the second displacement is sufficiently large or changes different feature combinations. Threshold sensitivity altered the class proportions, as expected, and is reported without selecting the most favorable threshold (Figure 3; the machine-readable path-class table in the code and data archive).

As a contextual control, whole-profile baseline RMSE also increased in negative-control and failed-reprogramming arms for all donors. These scalar controls were recovered from the archived all-profile pairwise-distance matrix; their treatment-arm values matched the new direct tensor calculations within 10^−7^. They restrict attribution of distance changes to reprogramming alone and are not causal treatment-effect estimates. Seven case-analysis checks verified shape and labels, finite values and masks, increment closure, nonnegative backtracking, its independent algebraic formula, exhaustive classification and reproduction of prior distances.

### 4.5 Do comparable paths occur in mock cultures?

We applied the same transition calculation to a parallel mock culture for each donor. Each pair shares its pretreatment sample. At the main tolerance (ε = 0.05), 8.23%, 8.12% and 5.53% of all treated sites departed and returned in O1, O2 and O3, respectively; mock rates were 4.74%, 4.74% and 3.74%. Among sites ending near baseline, the corresponding mock rates were 9.91%, 6.77% and 5.37%, compared with 15.72%, 13.95% and 9.75% in the treated series. The eligibility rule is the same in the two arms, but the sets of sites ending near baseline need not be identical (Supplementary Tables S2–S3).

A different question is whether sites that departed in both arms were more likely to return in the treated arm. On this shared set, the treated-minus-mock differences were +4.67, −6.52 and approximately zero percentage points across the three donors. Tightening ε to 0.02 also reversed the all-site contrast in O3 (−1.76 points). Removing sex-chromosome probes and probes flagged by the Zhou and Pidsley annotations [11,12] left 735,730 sites. At ε = 0.05 the all-site rates were 5.52–8.06% in treated and 3.59–4.60% in mock cultures (Supplementary Table S7). The observed return counts thus survive probe filtering, but their interpretation depends on the donor, the threshold and which sites enter the comparison.

### 4.6 What changes when visits or measurement rules are removed?

To measure the effect of fewer visits, we took ten untreated fibroblast culture series from six donors in GSE179847 [13]. For each series we kept the first and last visits and the visit closest to their elapsed-time midpoint. Among sites that departed from baseline and returned within 0.05 beta units on the denser schedule, the three-visit schedule missed 61.7% of those excursions. We first averaged the culture fractions within each donor and then weighted the six donors equally. Supplementary Tables S4, S5 and S8 give the counts, a stricter definition requiring consecutive departures, and a technical-replicate comparison in independent neutrophil series [14].

We also compared three implementations on 42 predefined questions with expected answers: 18 supported by the supplied measurements and 24 unsupported. EpiState-T and a pandas implementation enforcing the same rules answered all 18 supported questions and withheld answers to all 24 unsupported ones. A basic pandas script answered 10 of the 24 unsupported questions; it declined the other 14, once for a different reason from the prespecified one. Supplementary Table S6 gives the full accounting.

## 5 Discussion

### 5.1 What the representation adds

EpiState-T connects molecular arrays to the temporal and mechanistic claims being evaluated. In the executable model, it keeps current state, assumed neighbor influences, surviving records and reading-dependent transitions distinct. In the public-data case, it connects measured phases to explicit increments and path operators while marking unobserved mechanisms as unavailable. The distinction matters because the three observed phases can describe a return without measuring what caused it.

A longitudinal table can perform the same arithmetic if it preserves sample units, visits, features and treatment arms. The software comparison (§4.6) shows what is portable: rules about what a measurement can support. It does not establish a statistical advantage for tensor storage or estimate how often researchers make these errors in practice.

### 5.2 What the simulated reader establishes

Adding Read(H) resolves a concrete limitation of a retention-only example: changing memory availability can now change future states. Erasure and access blockade remove the effect, and content substitution changes its destination. Matched initial trajectories prevent an initial-state difference from explaining these model contrasts. The partial-retention result also exposes a condition under which recovery is not maintained.

Recovery toward an encoded checkpoint is nevertheless built into the rate law. Its successful execution demonstrates the behavior of that assumed controller, rather than discovering a biological restoration mechanism. The source-time query, payload accuracy and reading strength remain assumptions. Local historical reading is also distinct from historical information transfer among cells. Demonstrating a natural implementation would require measuring a candidate carrier and its content, perturbing access or content after formation, and testing recipient responses against present-regulation alternatives and appropriate functional outcomes.

### 5.3 What the three fibroblast samples per donor establish

The fibroblast case shows why historical representation matters even before a carrier is identified. Whole-profile methylation distances increased while thousands of loci showed observed departure and return. Conversely, a feature near baseline at the final sample could have remained near baseline or followed a different intermediate path. EpiState-T preserves the distinction and permits its magnitude to be calculated. The sampled-path result is the empirical application; it does not require fitting an unobserved H.

The reference is an adult pretreatment culture. Return toward that reference is not synonymous with age reversal, and departure is not synonymous with harm. The source study’s biological conclusions used other measurements and targets [1]. Our analysis addresses a narrower history question. Bulk populations also admit composition, differential survival, culture and technical effects; a returning average does not establish recovery by the same individual cells. Controls departing from baseline reinforce the need to keep observational descriptions separate from causal mechanisms.

The visit-removal comparison (§4.6) shows why “no departure observed at these visits” is more informative than “the site stayed at baseline.” Its miss fraction describes one sampling schedule in an independent dataset; it is not the probability that a particular Gill site had a hidden excursion.

### 5.4 Limitations and future comparisons

Three donors and three phases limit the application to descriptive analysis. The many CpGs improve feature coverage but do not increase the number of independent donors. Unequal sampling gaps, inherited preprocessing, all-profile QC selection, sensitivity to probe eligibility and the uncalibrated tolerance affect interpretation. The archived QC tensor permits precise reproduction of this analysis but does not constitute a fresh raw-data reconstruction. Additional observations within intervals would be needed to estimate more complete paths.

A future predictive test could compare models using only the current measurement with models that also use earlier measurements at matched measurement budgets, with preprocessing and model selection confined to training units. Such gains would still require controls for denoising and persistent unmeasured regulation.

Experiments with measurable retained content would be needed to test historical reading itself. By expressing these requirements in one representation, EpiState-T provides a route from a declared mechanism to observable comparisons without allowing simulated access to substitute for biological evidence.

## Supporting information

Supplementary File 2: Reproduction package (code and data)

## Data and code availability

The Gill fibroblast methylation data are in GEO (GSE165179; superseries GSE165180) [1,9]. The independent fibroblast sampling comparison uses GSE179847 [13]. The neutrophil technical-replicate checks use GSE247193 and GSE247195 [14]. The EpiState-T source repository is https://github.com/JadeSChoi/epistate-t (software v1.1.0, source at commit d8fbd30). Supplementary File 1 contains Figure S1 and Tables S1–S8. The complete reproduction package is Supplementary File 2 (EpiState-T_Paper_1_v1.3_Reproduction.zip). It contains the versioned code and data archive, including the processed focal tensor, sample identifiers and verification tests, plus the scripts for the supplement. Included counts and extracts reproduce the reported summaries offline; rerunning the full source-matrix analyses requires the deposited processed matrices. File hashes identify the versions used [10].

## Author contributions

Jade Sunghyeok Choi led conceptualization, methodology, software, formal analysis, visualization and original drafting. Jung In Choi contributed the original hypothesis about information distributed across cells, conceptual development and supervision. Both authors contributed to interpretation and manuscript review and editing.

## Ethics and AI assistance

No new biological intervention or participant recruitment was conducted. Public de-identified molecular data were analyzed. ChatGPT and Claude assisted earlier manuscript development; ChatGPT assisted this revision’s implementation, analysis, drafting and figure preparation. The human authors remain responsible for the scientific content and final approval.

## Relationship to the companion analysis

This methods paper and a separate analysis of RNA and DNA methylation reuse the same ten-day Gill cohort and methylation baseline distances. Here the empirical result concerns paths of individual CpG measurements across three sampled phases. The companion asks whether whole RNA and methylation profiles approach their donor’s starting profile and compares donor specificity and feature panels. These analyses use overlapping observations and are not independent biological replications.

