## Supplementary File 1: Figure S1 and Tables S1 to S8 for "EpiState-T for representing epigenetic states across time"

### A return in a culture measurement

Gill et al. [1] profiled populations of human fibroblasts before, during and after transient reprogramming. The panels compare a treated path with its matched mock path and show how often the same return criterion is met across the full array.

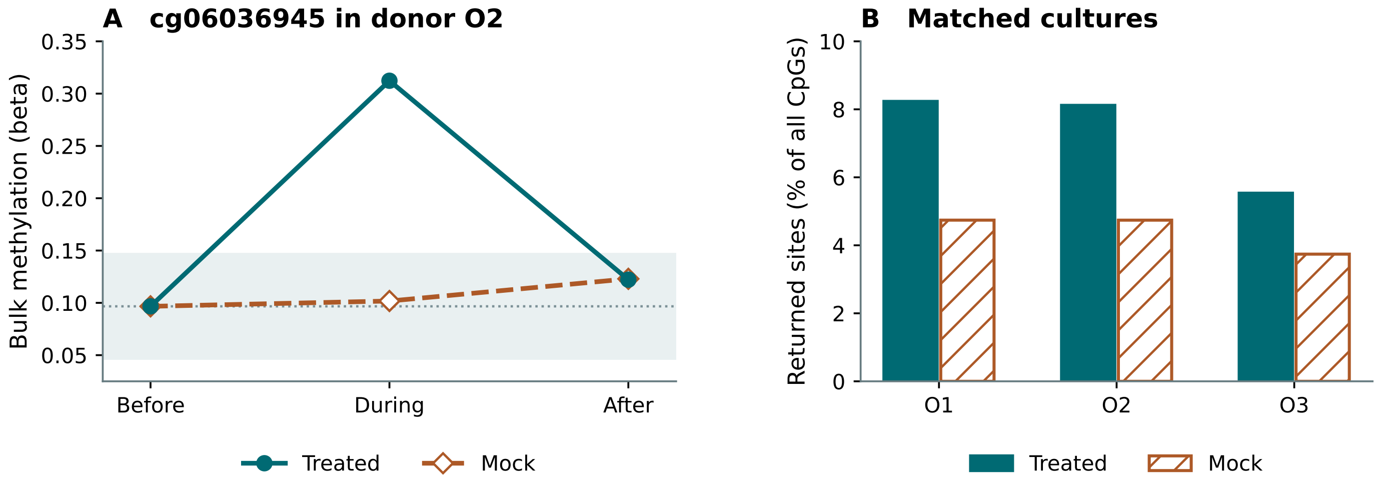


A. At cg06036945, a probe that passed both filters (Table S7), donor O2’s treated population departed from its baseline band and returned; the matched mock population never left that band at the sampled visits. Shading is ±0.05 beta units around the common baseline. Lines join ordered sampling phases, not equally spaced times.

B. At the same tolerance, the percentage of all 859,151 eligible sites classified as returning was higher in treated than mock cultures in each donor. Conditioning instead on sites that departed in both arms gives different contrasts (Table S3).

#### The two measured changes for panel A

| Arm | Before → during | During → after | Net change | Observed return |
|---|---|---|---|---|
| Treated | +0.216 | -0.190 | +0.026 | Yes |
| Mock | +0.005 | +0.021 | +0.026 | No departure |

Changes are beta units calculated before rounding. We selected cg06036945 from a fixed example panel by its treated path and probe annotation, before inspecting its mock outcome. This example illustrates the rule; panel B, not a selected probe, describes its frequency.

S1: simulation parameters; S2–S3: matched-arm comparisons; S4–S5: sampling and technical controls; S6: software checks; S7: probe masks; S8: confirmation across the full fibroblast array. Reference numbers follow the main text. The reference list is on page S9.

TABLE S1

### What record survival and reading strength change

Each entry is the final distance to each cell's own baseline after five model intervals. Rows specify the strength of reading (κ); columns specify the probability of preserving the checkpoint at each boundary (ρ). All 30 conditions use the same 100 seeds. Lower mean RMSE denotes closer states.

#### Mean final RMSE

| κ / ρ | 0 | 0.25 | 0.5 | 0.75 | 0.85 | 1 |
|---|---|---|---|---|---|---|
| 0 | 0.7443 | 0.7443 | 0.7443 | 0.7443 | 0.7443 | 0.7443 |
| 1 | 0.7443 | 0.7339 | 0.7029 | 0.6180 | 0.5386 | 0.3821 |
| 2 | 0.7443 | 0.7310 | 0.6898 | 0.5714 | 0.4764 | 0.2638 |
| 4 | 0.7443 | 0.7278 | 0.6766 | 0.5349 | 0.4281 | 0.1839 |
| 8 | 0.7443 | 0.7271 | 0.6702 | 0.5169 | 0.4086 | 0.1217 |

#### Monte Carlo standard error of each mean

| κ / ρ | 0 | 0.25 | 0.5 | 0.75 | 0.85 | 1 |
|---|---|---|---|---|---|---|
| 0 | 0.0052 | 0.0052 | 0.0052 | 0.0052 | 0.0052 | 0.0052 |
| 1 | 0.0052 | 0.0051 | 0.0052 | 0.0064 | 0.0076 | 0.0055 |
| 2 | 0.0052 | 0.0051 | 0.0059 | 0.0072 | 0.0097 | 0.0057 |
| 4 | 0.0052 | 0.0051 | 0.0060 | 0.0086 | 0.0103 | 0.0050 |
| 8 | 0.0052 | 0.0053 | 0.0061 | 0.0087 | 0.0108 | 0.0053 |

Approximate Monte Carlo intervals are mean ± 1.96 standard errors. With κ = 0 or ρ = 0, reading has no effect; these matching values use the same random seeds. At κ = 4 and ρ = 1 the mean is 0.1839 in this 100-seed sensitivity grid. The primary 500-seed estimate in the article is 0.1825.

**TABLE S2**

### The same return under two denominators

GSE165179: 859,151 quality-filtered CpGs per donor and arm. Let B, I and A be baseline, intermediate and after-treatment methylation. A return requires $|I - B| > \varepsilon$ and $|A - B| \le \varepsilon$. "Final-near" sites satisfy the latter condition in the specified arm. These are population measurements.

| ε | Donor | Treated returns | Mock returns | Treated % of all | Mock % of all | Treated % final-near | Mock % final-near |
|---|---|---|---|---|---|---|---|
| 0.02 | O1 | 90,267 | 56,048 | 10.51 | 6.52 | 40.06 | 31.64 |
| 0.02 | O2 | 99,932 | 90,020 | 11.63 | 10.48 | 30.93 | 24.32 |
| 0.02 | O3 | 71,393 | 86,503 | 8.31 | 10.07 | 22.42 | 23.05 |
| 0.05 | O1 | 70,733 | 40,728 | 8.23 | 4.74 | 15.72 | 9.91 |
| 0.05 | O2 | 69,734 | 40,724 | 8.12 | 4.74 | 13.95 | 6.77 |
| 0.05 | O3 | 47,543 | 32,137 | 5.53 | 3.74 | 9.75 | 5.37 |
| 0.10 | O1 | 29,962 | 12,201 | 3.49 | 1.42 | 5.21 | 1.90 |
| 0.10 | O2 | 28,585 | 6,619 | 3.33 | 0.77 | 4.55 | 0.88 |
| 0.10 | O3 | 21,418 | 4,653 | 2.49 | 0.54 | 3.39 | 0.63 |

Final-near denominators (treated / mock) at ε = 0.05: O1 449,837 / 410,847; O2 499,996 / 601,300; O3 487,663 / 598,375. The denominator uses the same rule in each arm but is not necessarily the same set of CpGs. The reproduction archive includes the table inputs for all tolerances.

#### Table S3. Restricting the comparison to sites that departed in both arms

| ε | Donor | Departed in both | Treated returns | Mock returns | Treated % | Mock % | Δ (pp) |
|---|---|---|---|---|---|---|---|
| 0.02 | O1 | 346,399 | 44,959 | 35,601 | 12.98 | 10.28 | +2.702 |
| 0.02 | O2 | 281,129 | 43,033 | 58,345 | 15.31 | 20.75 | -5.447 |
| 0.02 | O3 | 272,161 | 34,370 | 44,589 | 12.63 | 16.38 | -3.755 |
| 0.05 | O1 | 114,110 | 23,369 | 18,039 | 20.48 | 15.81 | +4.671 |
| 0.05 | O2 | 81,642 | 15,419 | 20,745 | 18.89 | 25.41 | -6.524 |
| 0.05 | O3 | 81,663 | 11,515 | 11,511 | 14.10 | 14.10 | +0.005 |
| 0.10 | O1 | 22,006 | 5,468 | 3,444 | 24.85 | 15.65 | +9.197 |
| 0.10 | O2 | 15,061 | 2,463 | 2,622 | 16.35 | 17.41 | -1.056 |
| 0.10 | O3 | 16,551 | 1,334 | 1,091 | 8.06 | 6.59 | +1.468 |

Δ = treated minus mock, in percentage points. The O3 difference of +0.005 points at ε = 0.05 is four returns among 81,663 common departing sites. This is a conditional descriptive comparison, not a treatment-effect adjustment. Source: Gill et al. [1].

TABLE S4

### Detours lost when only three visits are retained

Ten untreated fibroblast culture series from six donors (GSE179847 [13]) had 6–15 visits. A dense-schedule excursion ends within the baseline band after at least one observed departure. We retained both endpoints and the one measured visit nearest the elapsed-time midpoint. A “miss” is a known dense-schedule excursion no longer detectable in these three visits.

#### A. Tolerance sensitivity across all six donors

| ε | Dense returns | Retained on 3 visits | Missed | Missed % pooled | Missed % donor mean |
|---|---|---|---|---|---|
| 0.02 | 1,505,495 | 619,291 | 886,204 | 58.9 | 57.3 |
| 0.05 | 1,199,067 | 430,125 | 768,942 | 64.1 | 61.7 |
| 0.10 | 772,973 | 255,881 | 517,092 | 66.9 | 66.5 |

#### B. Per-donor results at ε = 0.05

| Donor | Culture series | Dense returns | Retained on 3 visits | Missed | Mean missed % within donor |
|---|---|---|---|---|---|
| HC1 | 3 | 390,752 | 108,823 | 281,929 | 71.9 |
| HC2 | 3 | 369,725 | 140,275 | 229,450 | 63.0 |
| HC3 | 1 | 113,686 | 48,402 | 65,284 | 57.4 |
| HC4 | 1 | 98,598 | 30,874 | 67,724 | 68.7 |
| HC5 | 1 | 120,119 | 47,053 | 73,066 | 60.8 |
| HC6 | 1 | 106,187 | 54,698 | 51,489 | 48.5 |

Counts pool culture–site paths. The donor-specific percentage first averages each culture’s missed fraction; it need not equal pooled missed / pooled dense in donors with multiple cultures. Each donor contributes equally to the headline estimate (61.7%). Pooling the counts instead gives 768,942 / 1,199,067 = 64.1%.

The dense schedule is an observed reference, not continuous ground truth. The denominator is dense-visible excursions; it is not all sites with a negative result after thinning. These estimates do not assign an individual CpG a probability of a hidden excursion.

TABLE S5

### Visit confirmation and technical replicates

These checks answer different questions. Part A asks whether two consecutive visits confirmed departure. Part B asks whether technical arrays at a single visit agreed. A second assay of one visit is not a second visit.

#### A. Fibroblasts: two originally consecutive departures on the same side

| Donor | Series | Dense, one departure | Dense, two consecutive | Four visits, confirmed | Missed %[1] |
|---|---|---|---|---|---|
| HC1 | 3 | 945 | 404 | 124 | 68.5 |
| HC2 | 3 | 891 | 390 | 174 | 63.7 |
| HC3 | 1 | 251 | 126 | 35 | 72.2 |
| HC4 | 1 | 233 | 57 | 15 | 73.7 |
| HC5 | 1 | 307 | 181 | 87 | 51.9 |
| HC6 | 1 | 265 | 131 | 78 | 40.5 |
| Pooled | 10 | 2,892 | 1,289 | 513 | 60.2 |

[1] Donor rows average per-culture fractions; the pooled row uses counts. This original 2,048-site check (20,480 site–culture paths; $\varepsilon = 0.05$) is reproduced from the full-array analysis in Table S8. Four visits retained both endpoints and the originally adjacent interior pair centered nearest the elapsed-time midpoint. On three visits the two-interior condition cannot be assessed. Two measured departures do not imply continuously displaced methylation between visits.

#### B. Neutrophils: visit mean and at least two agreeing technical arrays

| Donor series | ε | Dense returns | Three-visit returns | Missed | Missed % |
|---|---|---|---|---|---|
| GSE247193 | 0.02 | 167,135 | 48,700 | 118,435 | 70.9 |
| GSE247195 | 0.02 | 103,063 | 32,994 | 70,069 | 68.0 |
| GSE247193 | 0.05 | 31,802 | 6,575 | 25,227 | 79.3 |
| GSE247195 | 0.05 | 5,957 | 1,382 | 4,575 | 76.8 |
| GSE247193 | 0.10 | 1,967 | 300 | 1,667 | 84.7 |
| GSE247195 | 0.10 | 215 | 21 | 194 | 90.2 |

Independent donor series from Koncevičius et al. [14], with repeated technical arrays per visit. These losses are relative to dense observed paths, not technical false-positive rates. They cannot calibrate assay error in the Gill fibroblasts.

**TABLE S6**

### Conformance with the measurement contract

The primary test used 42 deliberately selected, independently labelled cases. An answer is unsupported if the requested biological or numerical interpretation cannot be justified by the supplied rows and metadata. A correct numerical operation can still give an unsupported answer.

#### A. Original test cases

| Workflow | Supported, correct | Unsupported, answered | Unsupported, refused with expected reason |
|---|---|---|---|
| Basic pandas | 18 / 18 | 10 / 24 | 13 / 24 |
| Checked pandas | 18 / 18 | 0 / 24 | 24 / 24 |
| EpiState-T | 18 / 18 | 0 / 24 | 24 / 24 |

Of 24 unsupported cases, basic pandas answered 10 and refused 14. Thirteen refusals gave the expected reason. In ob05 it correctly withheld an answer because the final value was missing; the specified reason was that the final visit was not observed.

#### B. Additional partial-date checks after the primary design

| Workflow | Permissible cases answered correctly | Contradictory date pairs answered |
|---|---|---|
| Basic pandas | 4 / 4 | 6 / 6 |
| Checked pandas, original rule | 4 / 4 | 6 / 6 |
| EpiState-T, original rule | 4 / 4 | 6 / 6 |
| Checked pandas, corrected | 4 / 4 | 0 / 6 |
| EpiState-T, corrected | 4 / 4 | 0 / 6 |

The correction checks all available dated pairs rather than waiting for a complete set. Corrected implementations refused the six contradictory cases and changed none of the original 42 responses. A sample swap carrying plausible but false labels remains undetectable from those labels alone. These tests do not measure how often researchers make errors.

**TABLE S7**

### Sensitivity to probe annotation

Gill data [1] at ε = 0.05. From the same 23-profile quality-filtered starting set, we retained autosomes (GRCh38 chromosomes 1–22), then excluded probes flagged by the Zhou EPIC general mask (Z) [11]. This excludes candidate mapping, sequence and common-variant problems, not all possible genotype effects. The last set additionally excludes candidate cross-reactive probes from the Pidsley list (P) [12].

| Probe set | Donor | Sites | Treated % all | Mock % all | Treated % final-near | Mock % final-near | Shared Δ (pp) |
|---|---|---|---|---|---|---|---|
| All QC sites | O1 | 859,151 | 8.23 | 4.74 | 15.72 | 9.91 | +4.671 |
| All QC sites | O2 | 859,151 | 8.12 | 4.74 | 13.95 | 6.77 | -6.524 |
| All QC sites | O3 | 859,151 | 5.53 | 3.74 | 9.75 | 5.37 | +0.005 |
| Autosomes | O1 | 840,007 | 8.23 | 4.62 | 15.58 | 9.62 | +4.968 |
| Autosomes | O2 | 840,007 | 8.11 | 4.71 | 13.84 | 6.69 | -6.713 |
| Autosomes | O3 | 840,007 | 5.50 | 3.68 | 9.63 | 5.26 | +0.010 |
| Autosomes + Z | O1 | 738,752 | 8.08 | 4.56 | 15.05 | 9.28 | +4.077 |
| Autosomes + Z | O2 | 738,752 | 7.98 | 4.61 | 13.54 | 6.50 | -6.945 |
| Autosomes + Z | O3 | 738,752 | 5.52 | 3.60 | 9.60 | 5.12 | +0.082 |
| Autosomes + Z + P | O1 | 735,730 | 8.06 | 4.56 | 15.02 | 9.27 | +4.023 |
| Autosomes + Z + P | O2 | 735,730 | 7.97 | 4.60 | 13.53 | 6.48 | -6.933 |
| Autosomes + Z + P | O3 | 735,730 | 5.52 | 3.59 | 9.60 | 5.12 | +0.095 |

Final-near percentages use each arm’s endpoint-near sites. Shared Δ compares treated minus mock returns among exactly the sites that had departed in both arms. Probe sets are nested; they are sensitivity checks, not independent replications. All ε = 0.02 and 0.10 results accompany the code.

#### A contrasting example excluded by the probe filters

cg24947637 lies on chromosome 17 and is flagged by Zhou’s MASK_snp5_GMAF1p and MASK_general. In O2 the common baseline was 0.753618; intermediate and final treated values were 0.852011 and 0.727630, versus 0.758482 and 0.745144 in mock. At ε = 0.05 the treated bulk signal meets the observed-return rule and the mock signal does not depart. The flag does not establish that the variant affected these donors. Figure S1A instead illustrates a probe retained by both filters.

**TABLE S8**

### Confirmation on the full fibroblast array

This analysis extends Table S5A to every eligible measured CpG in the ten Sturm cultures [13]. The tolerance, elapsed-time midpoint and confirmation rules are unchanged. A dense positive trajectory must have two originally consecutive, interior readings on the same side outside ±0.05, followed by a final reading near baseline. Three total visits are insufficient to test that definition.

#### A. Probe universe and available evidence

| Set | Eligible paths | Dense, one departure | Dense, two consecutive | Four visits, confirmed | Missed % pooled |
|---|---|---|---|---|---|
| 2,048 shared CpGs | 20,480 | 2,892 | 1,289 | 513 | 60.2 |
| All shared CpGs | 8,528,180 | 1,178,816 | 511,635 | 196,010 | 61.7 |
| All eligible in each culture | 8,601,914 | 1,199,067 | 521,296 | 199,529 | 61.7 |

#### B. All eligible sites per culture, aggregated by donor

| Donor | Series | Eligible paths | Dense, two consecutive | Four visits, confirmed | Missed | Missed % within donor |
|---|---|---|---|---|---|---|
| HC1 | 3 | 2,580,194 | 165,054 | 47,537 | 117,517 | 70.3 |
| HC2 | 3 | 2,580,145 | 158,748 | 68,692 | 90,056 | 61.9 |
| HC3 | 1 | 859,653 | 58,022 | 17,888 | 40,134 | 69.2 |
| HC4 | 1 | 861,289 | 23,385 | 7,368 | 16,017 | 68.5 |
| HC5 | 1 | 860,953 | 67,008 | 29,484 | 37,524 | 56.0 |
| HC6 | 1 | 859,680 | 49,079 | 28,560 | 20,519 | 41.8 |

Eligible paths count each culture × CpG once, not independent subjects. Four visits retain the two endpoints and one originally adjacent interior pair near the elapsed-time midpoint. The equal-donor missed fraction is 61.3% (range 41.8–70.3%). The 2,048-CpG subset was verified against the full recomputation. All results use the same ten cultures; expanding coverage is not replication in new donors.

**SOURCES**
